# Nephrobase Cell+: Multimodal Single-Cell Foundation Model for Decoding Kidney Biology

**DOI:** 10.1101/2025.09.30.679471

**Authors:** Chenyu Li, Elias Ziyadeh, Yash Sharma, Bernhard Dumoulin, Jonathan Levinsohn, Eunji Ha, Siyu Pan, Vishwanatha Rao, Madhav Subramaniyam, Mario Szegedy, Nancy Zhang, Katalin Susztak

## Abstract

**Background:** Large foundation models have revolutionized single-cell analysis, yet no kidney-specific model currently exists, and it remains unclear whether organ-focused models can outperform generalized models. The kidney’s complex cellular architecture and dynamic microenvironments further complicate integration of large-scale single-cell and spatial omics data, where current frameworks trained on limited datasets struggle to correct batch effects, capture cross-modality variation, and generalize across species.

**Methods:** We developed Nephrobase Cell+, the first kidney-focused large foundation model, pretrained on ~100 billion tokens from ~39.5 million single-cell and single-nucleus profiles across 4,319 samples, four mammalian species (human, mouse, rat, pig), and multiple assay modalities (scRNA-seq, snRNA-seq, snATAC-seq, spatial transcriptomics). Nephrobase Cell+ uses a transformer-based encoder-decoder architecture with gene-token cross-attention and a mixture-of-experts module for scalable representation learning.

**Results:** Nephrobase Cell+ sets a new benchmark for kidney single-cell analysis. It produces tightly clustered, biologically coherent embeddings in human and mouse kidneys, far surpassing previous foundation models such as Geneformer, scGPT, and UCE, as well as traditional methods such as PCA and autoencoders. It achieves the highest cluster concordance and batch-mixing scores, effectively removing donor/assay batch effects while preserving cell-type structure. Cross-species evaluation shows superior alignment of homologous cell types and >90% zero-shot annotation accuracy for major kidney lineages in both human and mouse. Even its 1B-parameter and 500M variants consistently outperform all existing models.

**Conclusions:** With organ-scale multimodal pretraining and a specialized transformer architecture, Nephrobase Cell+ delivers a unified, high-fidelity representation of kidney biology that is robust, cross-species transferable, and unmatched by current single-cell foundation models, offering a powerful resource for kidney genomics and disease research.

## Introduction

The advent of generative pretrained models has revolutionized artificial intelligence across diverse domains, from natural language processing to computer vision, by leveraging large-scale datasets and self-supervised learning frameworks to build versatile foundation models capable of generalizing across tasks^1, 2^. Inspired by these advances, the integration of deep learning with biological data has emerged as a transformative paradigm, particularly in single-cell genomics, where models such as scGPT^3^, Geneformer^4^ and scFoundation^5^ have demonstrated the power of pretraining on millions of cells to distill biological insights and enable transfer learning for downstream applications. By learning unified representations of genes, cells, and tissues, such models can capture biological context, mitigate technical noise, and extrapolate to unseen data^6, 7^. Despite significant progress in foundation models and omics technologies^8, 9^, kidney research remains constrained by fragmented analytical methods, limited scalability, and an incomplete understanding of cellular interactions in health and disease^10, 11^. Meanwhile, although general-purpose single-cell foundation models are rapidly emerging, organ-specific biology is still largely uncharted, and whether specialized models can outperform generalized ones in accuracy or interpretability remains an open question.

The kidney, a highly structured organ with diverse cell types and dynamic functional states, presents unique challenges and opportunities for such approaches. Kidney disease affects over 850 million individuals globally^12^, yet its molecular mechanisms remain poorly resolved due to the kidney’s cellular diversity and the multifactorial nature of pathologies such as fibrosis, inflammation, and metabolic dysregulation^13^. Single-cell RNA sequencing (scRNA-seq) and emerging multi-omic technologies have begun to unravel this complexity, revealing cell type-specific transcriptional programs, disease-associated states, and spatial organization patterns^14, 15^. Current computational methods in nephrology often rely on task-specific models trained on narrow datasets, limiting their ability to integrate multi-modal data, generalize across experimental conditions, or infer causal relationships. For instance, while tools like Seurat^16^ and Scanpy^17^ excel at clustering and dimensionality reduction, they lack the capacity to model hierarchical gene-regulatory networks or predict cellular responses to perturbations, which is a critical gap for therapeutic discovery. Furthermore, batch effects, sparse data, and inter-sample variability persist as major obstacles in large-scale kidney atlas initiatives like the Kidney Precision Medicine Project^18^.

Here, we present Nephrobase Cell+, the largest single-cell foundational model to date and the first kidney-specific model. Instead of fine-tuning an existing model, we pretrained a foundational model for kidney biology from scratch on over 100 billion tokens and 39 million cells and nuclei derived from more than samples spanning human and murine kidneys across health, developmental, and disease states. Nephrobase Cell+ integrates scRNA-seq, single-nucleus RNA-seq (snRNA-seq), and spatially resolved transcriptomic data, employing a masked generative pretraining strategy adapted from large language models to learn robust representations of kidney cell identities, gene regulatory networks, and microenvironmental crosstalk. By unifying data from diverse technologies, donors, and species, the model addresses key limitations of existing methods, including batch effects, modality-specific biases, and sparse gene coverage.

## Results

### Dataset composition and sampling breadth

To train Nephrobase Cell+, we assembled a large, diverse multimodal kidney atlas comprising 4,319 samples and ~39.5 million single-cell/single-nucleus profiles (Fig. 1). The combined dataset spans four mammalian species and multiple biological contexts and assays. Breakdown by species is as follows: Homo sapiens: 25.2 million cells, Mus musculus: 12.5 million cells, Rattus norvegicus: 1.4 million cells, and Sus scrofa: 293,000 cells. Across tissues and sample sources, approximately ~30 million of the profiles derive from kidney tissue, with an additional ~10 million profiles arising from immune and peripheral sources, reflecting the inclusion of peripheral blood mononuclear cells and immune-enriched samples. Assay composition reveals broad multimodality: single-cell RNA-seq (scRNA-seq) comprises the largest fraction of profiles (~48.7%), followed by snRNA-seq (~25.4%). Spatial transcriptomics modalities were represented by COSMx (~7.7%), and Xenium runs (~5.5%). Single-nucleus ATAC-seq (snATAC-seq) contributed roughly ~6.2% of assays. The predominance of RNA-based assays combined with substantial spatial and chromatin accessibility data provided a strong multimodal training signal for cross-assay representation learning.This scale and diversity enabled robust learning of conserved cellular programs while also providing substantial representation of species- and assay-specific variation that the model was trained to mitigate.

**Figure 1.**
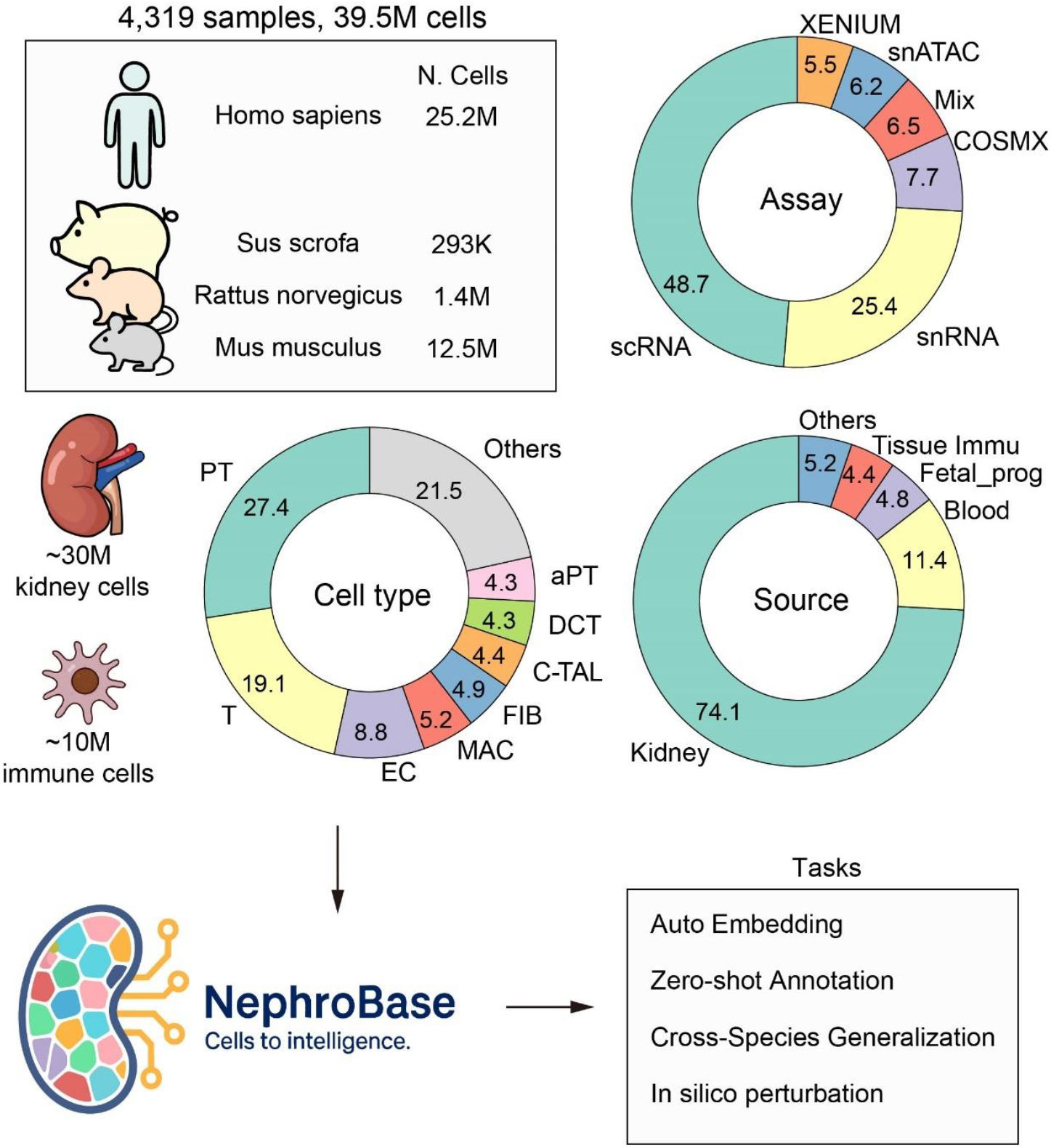
Composition of the Nephrobase Cell+ training dataset. The Nephrobase Cell+ atlas integrates 4,319 samples encompassing ~39.5 million single-cell and single-nucleus profiles. Top left: Distribution of cells across four mammalian species: Homo sapiens (25.2M), Mus musculus (12.5M), Rattus norvegicus (1.4M), and Sus scrofa (0.293M). Bottom left: Approximate tissue origin of the dataset, including ~30M kidney-derived cells and ~10M immune cells. The cell-type composition shows strong representation of proximal tubule (PT, 27.4%), T cells (19.1%), endothelial cells (EC, 8.8%), macrophages (MAC, 5.2%), fibroblasts (FIB, 4.9%), cortical thick ascending limb (C-TAL, 4.3%), distal convoluted tubule (DCT, 4.3%), and atrophic proximal tubule (aPT, 4.3%), with the remainder categorized as “others” (21.5%). Top right: Assay composition, highlighting contributions from scRNA-seq (48.7%), snRNA-seq (25.4%), COSMx (7.7%), Xenium (5.5%), snATAC-seq (6.2%), and mixed modalities (6.5%). Bottom right: Sample source composition, showing that most profiles are from kidney tissue (74.1%), with additional contributions from blood (11.4%), fetal/progenitor samples (4.8%), tissue-enriched immune fractions (4.4%), and other sources (5.2%). Together, these distributions illustrate the multimodal, multispecies, and multicontext diversity of the Nephrobase Cell+ training dataset.

### Overview of model architecture and training strategy

We developed Nephrobase Cell+, a transformer-based encoder-decoder framework augmented with modular components tailored to single-cell/single-nucleus and spatial transcriptomic data integration. The model ingests a cell-gene matrix and produces both reconstructed gene expression distributions and cell type predictions in a unified architecture. As illustrated in Fig. 2A: an initial gene tokenization step converts gene identities, observed expression and optional metadata into token embeddings by cross attention; and the resulting tokens are processed with self-attention in the encoder and cross-attention in the decoder. The reconstruction head optimizes a Zero-Inflated Negative Binomial likelihood to capture count overdispersion and excess zeros typical of single-cell data, while a classification head is trained with a focal loss to address class imbalance and emphasize difficult examples (Fig. 2B).

**Figure 2.**
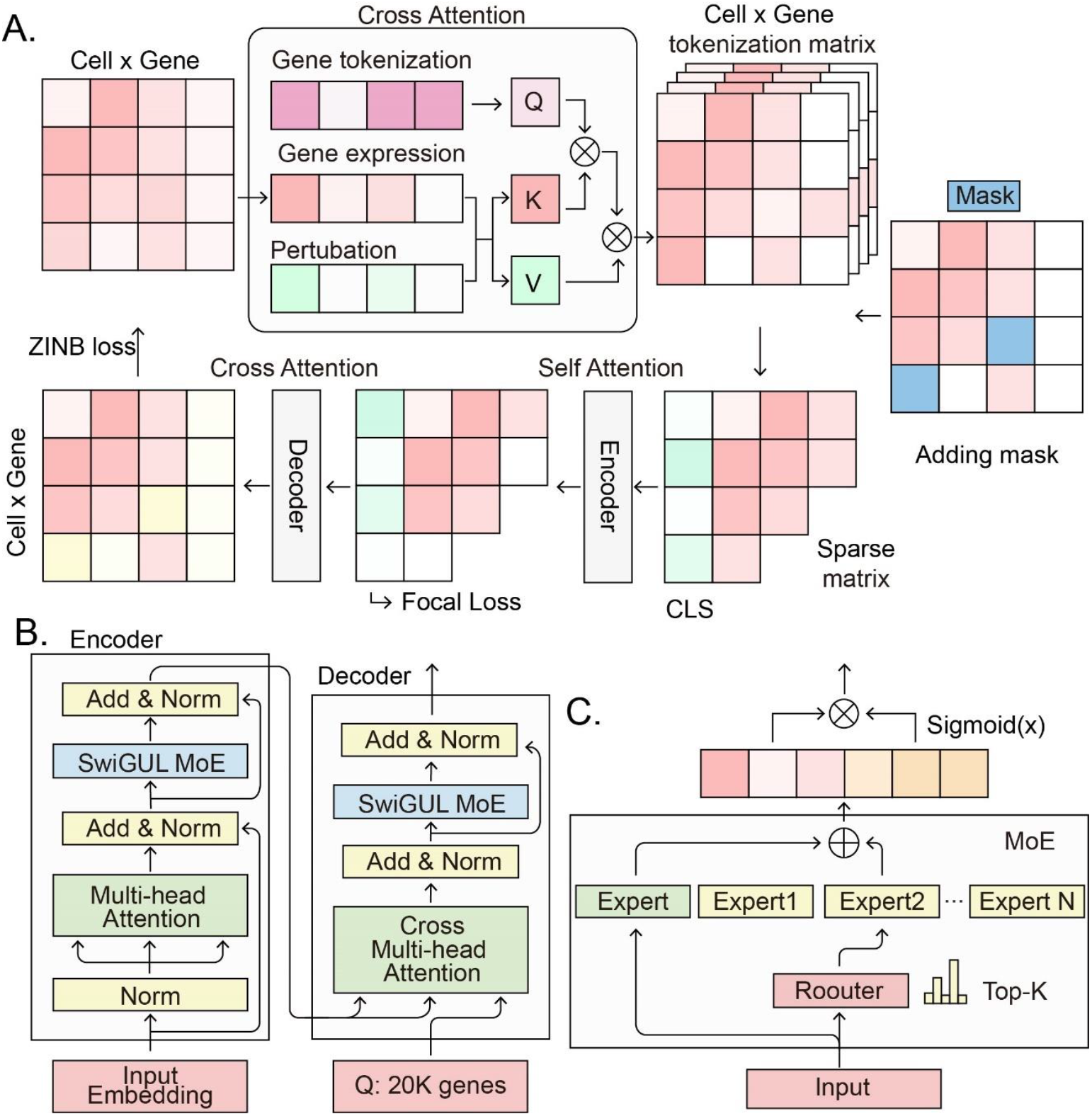
Nephrobase Cell+ model architecture and training strategy. (A) Overview of the encoder-decoder framework. The model ingests a cell-by-gene matrix, with each gene tokenized using its identity, normalized expression value, and optional perturbation metadata. Tokens are embedded via cross-attention to generate a cell × gene tokenization matrix, with masking applied to subsets of inputs. The encoder applies self-attention, while the decoder uses cross-attention to reconstruct expression profiles. Outputs are optimized using a Zero-Inflated Negative Binomial (ZINB) loss for count reconstruction and a focal loss for supervised cell-type classification. Detailed transformer block design. Both encoder and decoder stacks include normalization layers, multi-head attention modules, and SwiGUL Mixture-of-Experts (MoE) layers, with cross multi-head attention connecting the decoder to the encoder. Input embeddings represent up to 20,000 genes per cell. (C) Structure of the Mixture-of-Experts (MoE) module. Each input is routed to a subset of specialized experts using top-k gating, with outputs combined through weighted summation. A shared expert and sigmoid activation further stabilize and generalize representation learning. Together, these components allow Nephrobase Cell+ to learn robust, assay-invariant embeddings of kidney cell states from large-scale multi-modal data.

A Mixture-of-Experts module (Fig. 2C) expands model capacity via sparse top-k routing to a set of specialized experts and includes a shared-expert extension to capture global transformations; a load-balancing auxiliary loss encourages even expert utilization. To prevent representational collapse and to shape the embedding geometry, we combined an Elastic Cell Similarity regularizer that enforces a target level of dissimilarity between cell embeddings with a supervised contrastive loss that pulls together cells of the same annotation and pushes apart differently labelled cells. Finally, adversarial discriminators with a gradient reversal layer were employed to remove assay and batch signals from learned features, producing more assay-invariant representations for downstream classification and reconstruction. We trained two Nephrobase Cell+ models, with approximately 1 billion (1B) and 500 million (500M) parameters, respectively (Table 1).

**Table 1.** Model Architecture and Training Hyperparameters of Nephrobase Cell+ Variants*.

|  | Nephrobase Cell+<br>1B | Nephrobase Cell+<br>500M |
| --- | --- | --- |
| embed_d | 1024 | 768 |
| batch_size | 90 | 136 |
| enc_nheads | 8 | 8 |
| dec_nheads | 8 | 8 |
| n_fuse_attention | 1 | 1 |
| n_encoder | 6 | 5 |
| n_decoder | 1 | 1 |
| enc_mlp_ratio | 4 | 4 |
| dec_mlp_ratio | 4 | 4 |
| num_classes | 31 | 31 |
| Training strategy | FSDP | FSDP |
| Parameter | ~500M | ~1B |

\*embed\_d, embedding dimension; enc\_nheads, number of encoder attention heads; dec\_nheads, number of decoder attention heads; n\_fuse\_attention, number of fusion attention layers; n\_encoder, number of encoder layers; n\_decoder, number of decoder layers; enc\_mlp\_ratio, encoder multilayer-perceptron ratio; dec\_mlp\_ratio, decoder multilayer-perceptron ratio; FSDP, Fully Sharded Data Parallel.

### Integrated Embedding Visualization and Clustering

To assess how Nephrobase Cell+ representations capture kdieny cell identities, we compared its learned embeddings to those from baseline methods on held-out human and mouse kidney single-cell RNA-seq data. We applied UCE, Principal Component Analysis (PCA), and an autoencoder on each model’s latent space and visualized the results with UMAP (Figure 3). We selected datasets from two species, neither of which had been used for training. One dataset comes from KPMP’s 2025 Q2 data, and the other comes from a mouse developmental model. In both species, Nephrobase Cell+ produced embeddings yield clear, compact clusters corresponding to known kidney cell. For instance, in human data (Fig. 3A), Nephrobase Cell+ 1B-parameter embeddings separate proximal tubule (PT) and Thick Ascending Limb (TAL) into distinct groups, whereas Geneformer, UCE and scGPT produce more diffuse or mixed clusters. Similarly, in mouse (Fig. 3B) Nephrobase Cell+ distinguishes PT, TAL, immune, podocytes, and progenitor cells more cleanly than competing models.

**Figure 3.**
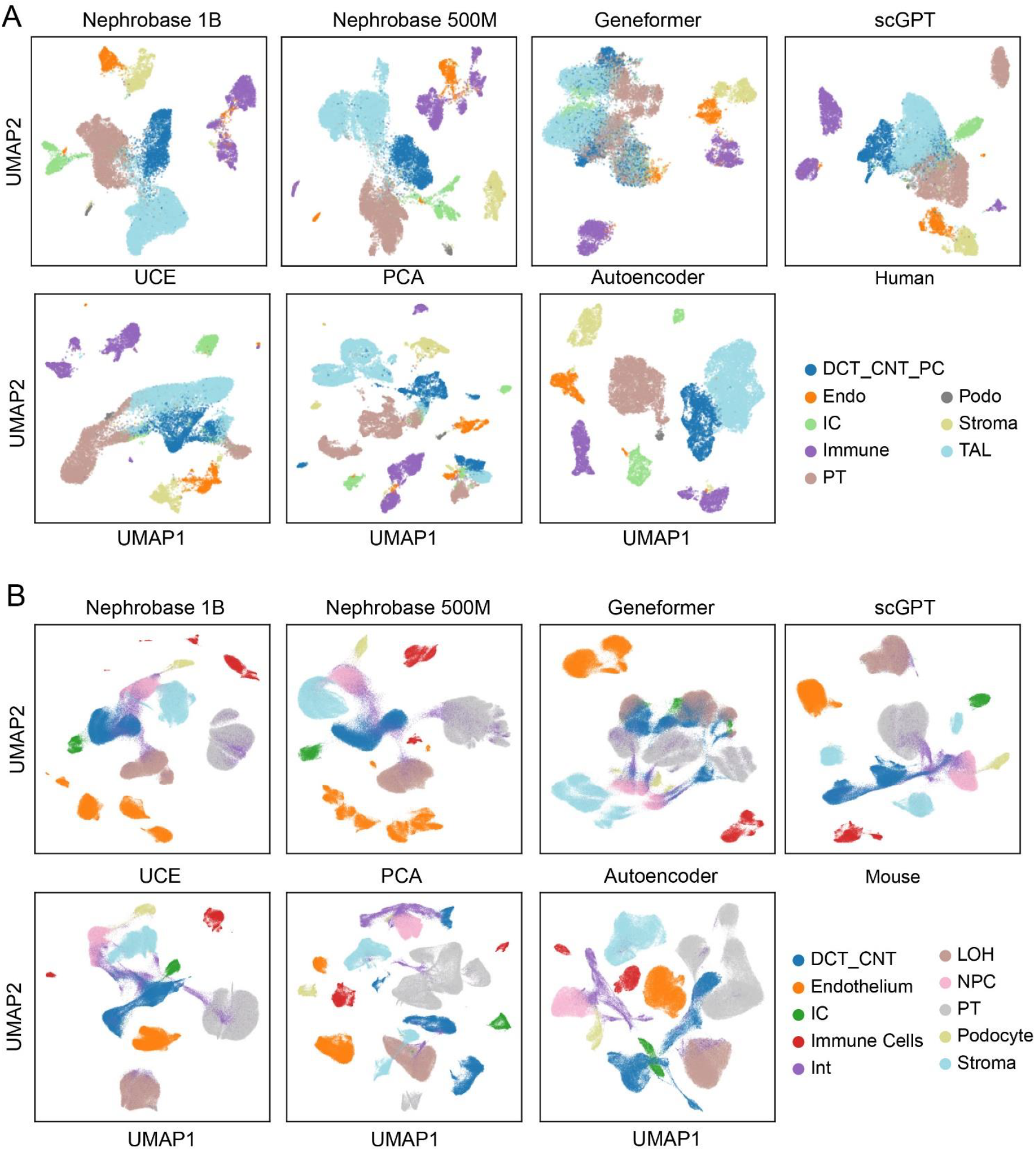
Zero-shot benchmarking of embedding of single-nucleus transcriptomic data analysis methods using for human and mouse datasets. (A) UMAP projections of human kidney data from various single-cell RNA-seq analysis methods: Nephrobase Cell+ 1B, Nephrobase Cell+ 500M, Genformer, and scGPT. Each method is shown with clustering using Uniformed Cluster Embedding (UCE), Principal Component Analysis (PCA), and Autoencoder dimensionality reduction techniques. Proximal Tubule (PT) (cyan), Stroma (yellow), and TAL (light blue). (B) UMAP projections of mouse kidney data following the same methods and dimensionality reduction techniques as in (A). DCT_CNT_PC (Distal Convoluted Tubule and Connecting Tubule Principal Cells), Endo (Endothelium), IC (Intercalated Cells), Immune (Immune Cells), Podo (Podocytes), PT (Proximal Tubule), Stroma (Stromal Cells), TAL (Thick Ascending Limb), LOH (Loop of Henle), NPC (Nephron Progenitor Cells), Int (Interstitial Cel

These qualitative observations are supported by quantitative clustering metrics^19^: In human data, both Nephrobase Cell+ variants achieved the highest cell-type isolation scores: KMeans ARI is 0.82 for the 1B model (versus 0.40 for scGPT, 0.55 for autoencoder, and only 0.22 for Geneformer), and NMI is 0.78 (vs 0.48 scGPT). The Silhouette score for Nephrobase Cell+ (~0.68) also exceeds that of competitors. Importantly, Nephrobase Cell+ fully integrates cells from different samples: the cLISI score is 1.00 (perfect batch mixing) for both 1B and 500M models, while iLISI (label-agnostic mixing) is very low (~0.17, 0.18 in human), indicating minimal batch-driven separation. Batch-correction tests (kBET) are correspondingly low (0.25-0.28 for Nephrobase Cell+ vs 0.09 for scGPT, where lower is better). As a result, Nephrobase Cell+ attains the highest overall integration score (Total = 0.71 for 1B, 0.70 for 500M) among all evaluated methods (Table 2, bottom row). In mouse, similar trends are observed: Nephrobase Cell+ yields ARI ~0.70 (higher than all baselines) and cLISI=1.0, with Total scores of 0.69-0.68, again outperforming UCE, PCA, and scGPT. Together, Figure 3 demonstrates that Nephrobase Cell+ integrates multi-assay kidney data into a latent space that strongly reflects underlying biology, producing visually and quantitatively superior clustering of cell types compared to existing dimensionality-reduction or single-cell model baselines.

**Table 2.** Batch effect removing benchmarking Nephrobase Cell+ against existing foundational and dimensionality reduction models in human and mouse datasets.*

|  |  | Nephrobase | Nephrobase | Autoencoder | Geneformer | UCE | PCA | scGPT |
| --- | --- | --- | --- | --- | --- | --- | --- | --- |
| Metrics |  | Cell+<br>1B | Cell+<br>500M |  |  |  |  |  |
| Human | Isolated labels | 0.76 | 0.77 | 0.75 | 0.58 | 0.66 | **0.89** | 0.62 |
|  | KMeans NMI | **0.78** | 0.76 | 0.72 | 0.37 | 0.63 | 0.54 | 0.48 |
|  | KMeans ARI | **0.82** | 0.67 | 0.55 | 0.22 | 0.48 | 0.4 | 0.3 |
|  | Silhouette label | **0.68** | 0.67 | 0.62 | 0.52 | 0.6 | 0.58 | 0.55 |
|  | cLISI | **1** | **1** | **1** | 0.97 | **1** | **1** | **1** |
|  | BRAS | 0.74 | **0.77** | 0.79 | 0.77 | 0.72 | 0.28 | 0.71 |
|  | iLISI | 0.17 | **0.18** | 0.13 | 0.12 | 0.06 | 0.01 | 0.09 |
|  | KBET | 0.25 | **0.28** | 0.1 | 0.12 | 0.07 | 0.09 | 0.1 |
|  | Graph connectivity | 0.94 | 0.93 | **0.98** | 0.8 | 0.84 | 0.71 | 0.88 |
|  | PCR comparison | 0.67 | 0.73 | **0.84** | 0.44 | 0.38 | 0.13 | 0.36 |
|  | Batch correction | 0.55 | **0.58** | 0.57 | 0.45 | 0.41 | 0.24 | 0.43 |
|  | Bio conservation | **0.81** | 0.77 | 0.73 | 0.54 | 0.67 | 0.68 | 0.59 |
|  | Total | **0.71** | 0.7 | 0.67 | 0.5 | 0.57 | 0.51 | 0.53 |
| Mouse | Isolated labels | 0.64 | 0.62 | 0.77 | 0.58 | 0.65 | **0.87** | 0.66 |
|  | KMeans NMI | **0.79** | **0.79** | **0.79** | 0.6 | 0.75 | 0.69 | 0.79 |
|  | KMeans ARI | **0.7** | **0.68** | 0.65 | 0.41 | 0.6 | 0.53 | 0.65 |
|  | Silhouette label | **0.69** | 0.66 | 0.63 | 0.55 | **0.67** | 0.55 | 0.65 |
|  | cLISI | 1 | 1 | 1 | 1 | 1 | 1 | 1 |
|  | BRAS | 0.88 | 0.88 | 0.83 | **0.89** | 0.86 | 0.55 | 0.86 |
|  | iLISI | **0.21** | **0.21** | 0.14 | 0.17 | 0.15 | 0.16 | 0.16 |
|  | KBET | **0.44** | 0.48 | 0.17 | 0.31 | 0.18 | 0.18 | 0.31 |
|  | Graph connectivity | 0.93 | 0.92 | **0.99** | 0.89 | 0.86 | 0.82 | 0.88 |
|  | PCR comparison | 0.24 | **0.51** | 0.15 | 0.02 | 0 | 0 | 0 |
|  | Batch correction | 0.54 | **0.6** | 0.45 | 0.46 | 0.41 | 0.34 | 0.44 |
|  | Bio conservation | 0.76 | 0.75 | **0.77** | 0.63 | 0.73 | 0.73 | 0.75 |
|  | Total | 0.67 | **0.69** | 0.64 | 0.56 | 0.6 | 0.57 | 0.63 |

\*NMI, normalized mutual information; ARI, adjusted Rand index; cLISI, cell-type Local Inverse Simpson's Index; iLISI, integration Local Inverse Simpson's Index; BRAS, batch removal average silhouette; KBET, k-nearest neighbor batch effect test; PCR, principal component regression.

### Cross-Species Generalization

Next, we evaluated Nephrobase Cell+’s ability to generalize across species. We projected a mixed human-mouse kidney dataset into each model’s embedding space and compared the resulting UMAP visualizations (Figure 4). Each row of Figure 4 corresponds to a different model: Nephrobase Cell+ (1B and 500M), Geneformer, scGPT, and UCE applied to raw data. The columns show species origin, manual cell-type annotations, and zero-shot model cell-type predictions. Ideally, human and mouse cells of the same type should cluster together and receive the same predicted labels. Nephrobase Cell+ achieves this ideal alignment: human and mouse proximal tubule, TAL, DCT/CNT, IC, podocytes, stromal, endothelial, and immune cells form overlapping clusters and Nephrobase Cell+’s predicted labels match the manual annotations for both species. In contrast, Geneformer and UCE embeddings show pronounced species segregation and poorer agreement with expert labels. scGPT performs reasonably but still shows more mixing errors than Nephrobase Cell+. These patterns indicate that Nephrobase Cell+ has learned species-invariant features of kidney cells. The zero-shot predictions from Nephrobase Cell+ recapitulate the annotated taxonomy with high fidelity, even though the model was not fine-tuned on this held-out data. Overall, Figure 4 illustrates that Nephrobase Cell+ produces an integrated cross-species embedding in which homologous cell types are co-localized and correctly identified, whereas other models exhibit weaker cross-species alignment and more frequent misclassification.

**Figure 4.**
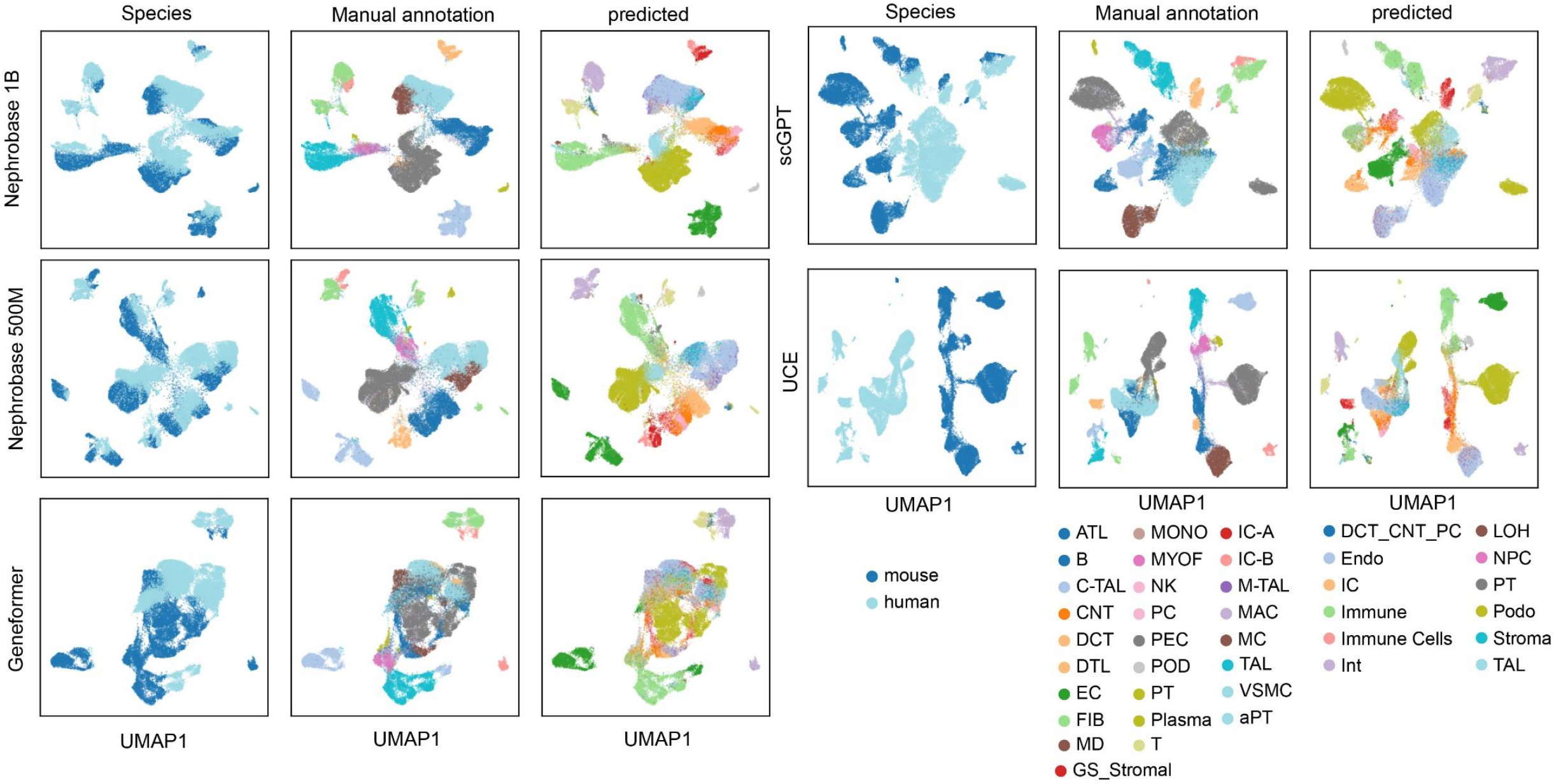
Zero-shot cross-species benchmarking of foundation models for kidney single--nucleus transcriptomics. UMAP visualizations of human and mouse kidney single-cell transcriptomic data comparing species labels, manual expert annotations, and model-predicted cell types across multiple foundation models for Nephrobase Cell+ models (1B and 500M parameters),Geneformer, scGPT and UCE. Each row corresponds to a distinct model, with columns showing (i) species distribution (human, light blue; mouse, dark blue), (ii) manual cell type annotations, and (iii) zero-shot model predictions. Major kidney epithelial, stromal, endothelial, and immune cell types are highlighted, including proximal tubule (PT), thick ascending limb (TAL), distal convoluted tubule/connecting tubule (DCT/CNT), intercalated cells (IC), podocytes (PODO), stromal cells, endothelial cells (Endo), immune cells, and nephron progenitors (NPC). Predictions from Nephrobase Cell+ and scGPT more closely recapitulate expert manual annotations compared to Geneformer and UCE, demonstrating improved cross-species generalizability and fine-grained nephron cell type resolution.

For the metrics^19^, Nephrobase Cell+ again achieves superior scores: the 500M model attains NMI = 0.75 and ARI = 0.72 on KMeans clustering (versus 0.44/0.22 for Geneformer, 0.62/0.43 for scGPT). The larger 1B model has nearly identical results (NMI = 0.73, ARI = 0.57). Both models reach perfect label mixing across species (cLISI = 1.00) and very low iLISI (0.01), indicating that human and mouse cells are indistinguishable in Nephrobase Cell+’s latent space. Graph connectivity and PCR values are also highest for Nephrobase Cell+ (Graph conn. ~0.94, PCR ~0.94), reflecting well-connected and unbiased embeddings. The overall integration score for Nephrobase Cell+ 500M is 0.63 (1B: 0.61), substantially above Geneformer (0.50) and UCE (0.51, Table 3).

**Table 3.** Cross-species benchmarking Nephrobase Cell+ against existing foundational and dimensionality reduction models in human and mouse datasets.*

| Metrics | Nephrobase Cell+ | Nephrobase Cell+ | Geneformer | UCE | scGPT |
| --- | --- | --- | --- | --- | --- |
|  | 1B | 500M |  |  |  |
| Isolated labels | 0.54 | 0.54 | 0.56 | **0.64** | 0.56 |
| KMeans NMI | 0.73 | **0.75** | 0.44 | 0.7 | 0.62 |
| KMeans ARI | 0.57 | **0.72** | 0.22 | 0.53 | 0.43 |
| Silhouette label | **0.61** | 0.6 | 0.51 | 0.6 | 0.57 |
| cLISI | 1 | 1 | 0.99 | 1 | 1 |
| BRAS | 0.6 | 0.62 | **0.69** | 0.38 | 0.64 |
| iLISI | **0.01** | **0.01** | 0 | 0 | 0 |
| KBET | **0.04** | 0.03 | 0.02 | 0 | 0.01 |
| Graph connectivity | **0.88** | 0.84 | 0.84 | 0.72 | 0.85 |
| PCR comparison | **0.94** | 0.94 | 0.61 | 0.04 | 0.8 |
| Batch correction | **0.49** | 0.49 | 0.43 | 0.23 | 0.46 |
| Bio conservation | 0.69 | **0.72** | 0.54 | 0.69 | 0.64 |
| Total | 0.61 | **0.63** | 0.5 | 0.51 | 0.57 |

\*NMI, normalized mutual information; ARI, adjusted Rand index; cLISI, cell-type Local Inverse Simpson's Index; iLISI, integration Local Inverse Simpson's Index; BRAS, batch removal average silhouette; KBET, k-nearest neighbor batch effect test; PCR, principal component regression.

### Zero-Shot Cell-Type Annotation

To quantify Nephrobase Cell+’s classification performance, we performed zero-shot cell-type prediction on held-out human and mouse test sets and compared to manual curation. Figure 5 shows confusion matricesand Sankey diagrams for human and mouse data. In each confusion matrix, rows represent manual expert labels and columns represent Nephrobase Cell+’s predicted labels. The darkest diagonal elements indicate the fraction of cells correctly identified. In human kidney (Fig. 5A), Nephrobase Cell+ correctly annotates the majority of cells in all major nephron lineages: for example, over 90% of PT cells are predicted as PT, and similarly high agreement is seen for TAL, DCT, CNT, intercalated (IC), and podocyte classes. Minor misclassifications mostly occur between closely related subtypes (e.g. distal tubule vs connecting tubule). The human Sankey diagram (Fig. 5B) visualizes these relationships: thick flows along the diagonal demonstrate that manual and predicted labels align, with only a few thin off-diagonal flows. Mouse results are analogous (Fig. 5C,D): Nephrobase Cell+ achieves high concordance for nephron progenitors, PT, TAL, podocytes, stromal, endothelial, and immune cells. Across both species, the overall accuracy of zero-shot annotation exceeds that of simpler methods (not shown) and matches expert-level curation for major cell groups. These results confirm that the Nephrobase Cell+ embeddings and classification head generalize to new data: the model effectively transfers learned cell-type signatures without retraining, yielding robust annotation consistent with manual standards.

**Figure 5.**
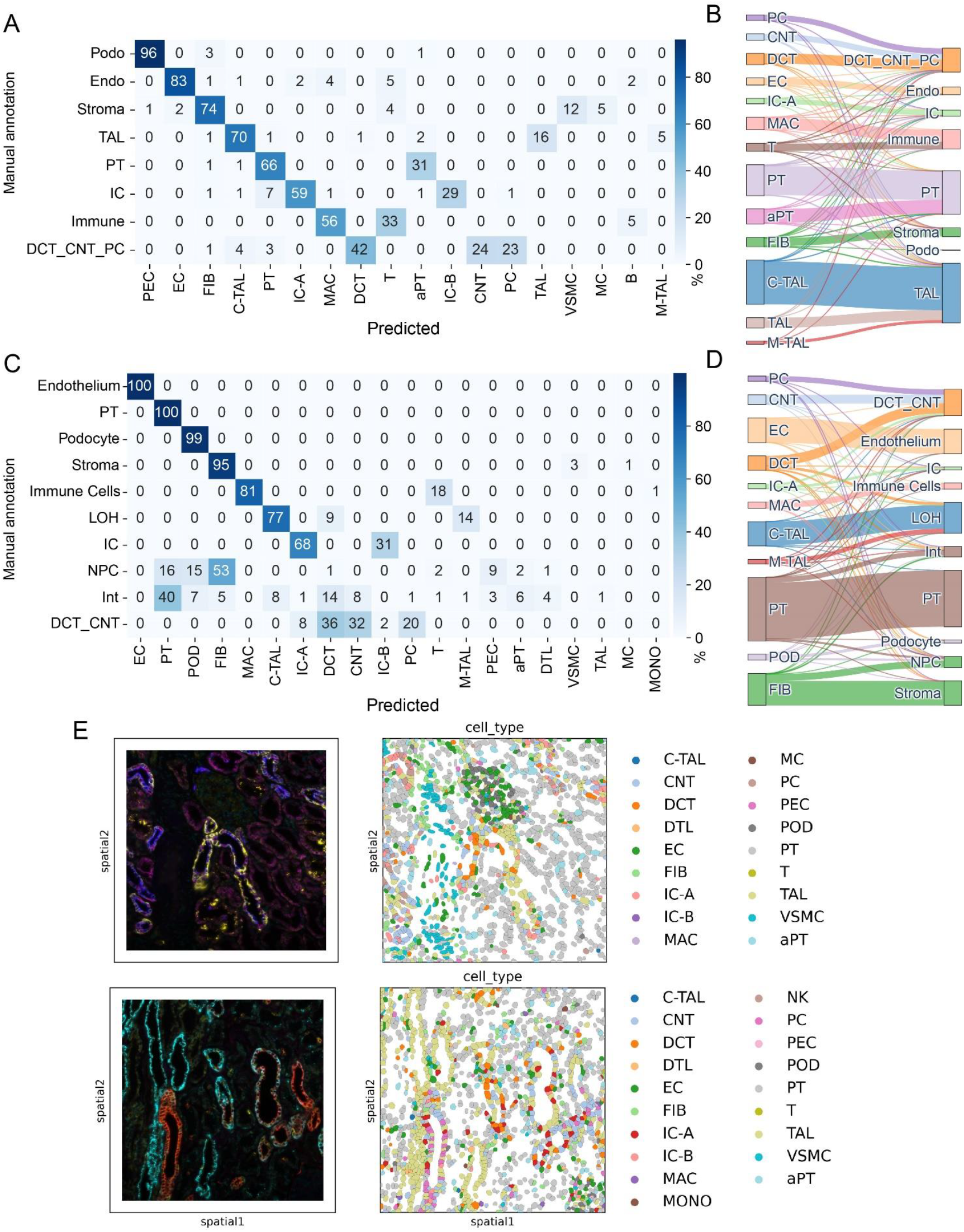
Zero-shot benchmarking Nephrobase Cell+ cell type annotation against manual curation. Confusion matrices showing agreement between manual annotations (rows) and Nephrobase Cell+ predictions (columns) across major kidney cell types in (A) human and (C) mouse datasets. The percentage of cells correctly assigned to each category is indicated by the color intensity, with darker shades reflecting higher concordance. (B, D) Sankey diagrams illustrating the mapping between manual annotations and Nephrobase Cell+ predictions for the same datasets. (E) an example should the predict cell type for spatial transcriptome.Major nephron epithelial lineages (PT, TAL, DCT, CNT, IC, Podocytes) as well as stromal, endothelial, and immune compartments are shown. The width of each connection is proportional to the number of cells assigned. Together, these analyses demonstrate that Nephrobase Cell+ achieves high concordance with expert manual curation while capturing fine-grained nephron subtypes.

### In silico perturbation

We performed a simple perturbation by randomly sampling 5,000 cells and doubling the expression of each target gene, then ranked genes by differential expression and ran gene set enrichment analysis to identify perturbed biological programs (Fig 6). Perturbation of CCL2 produced a clear proinflammatory and chemotactic signature with significant enrichment of lymphocyte activation, leukocyte cell adhesion, macrophage migration and eosinophil chemotaxis, together with ion transmembrane transport terms, indicating concurrent effects on immune recruitment and membrane transport. Perturbation of VCAM1 similarly enriched adhesion and immune activation processes, including T cell proliferation, leukocyte adhesion to vascular endothelium and neutrophil chemotaxis, and also highlighted vascular endothelial migration and cellular respiration pathways such as aerobic respiration and hydrogen peroxide catabolism, consistent with a link between endothelial remodeling and metabolic adaptation. GDF15 perturbation shifted the transcriptional profile toward growth regulation and signaling, with enrichment of negative regulation of developmental growth, fructose metabolic process, receptor tyrosine kinase signaling and MAPK cascade terms, while also showing leukocyte aggregation and neutrophil migration signatures. Perturbation of SOX4 produced prominent developmental and bioenergetic responses, with enrichment of organ morphogenesis terms including kidney morphogenesis and valve morphogenesis, programmed cell death, proton transport and oxidative phosphorylation, suggesting simultaneous impacts on developmental programs and mitochondrial function. These results indicate that simple twofold upregulation of individual genes elicits distinct and biologically coherent transcriptional responses, with CCL2 and VCAM1 predominantly driving immune adhesion and chemotaxis, GDF15 modulating growth control and MAPK signaling with immune aggregation features, and SOX4 strongly affecting developmental processes and cellular energy metabolism.

**Figure 6.**
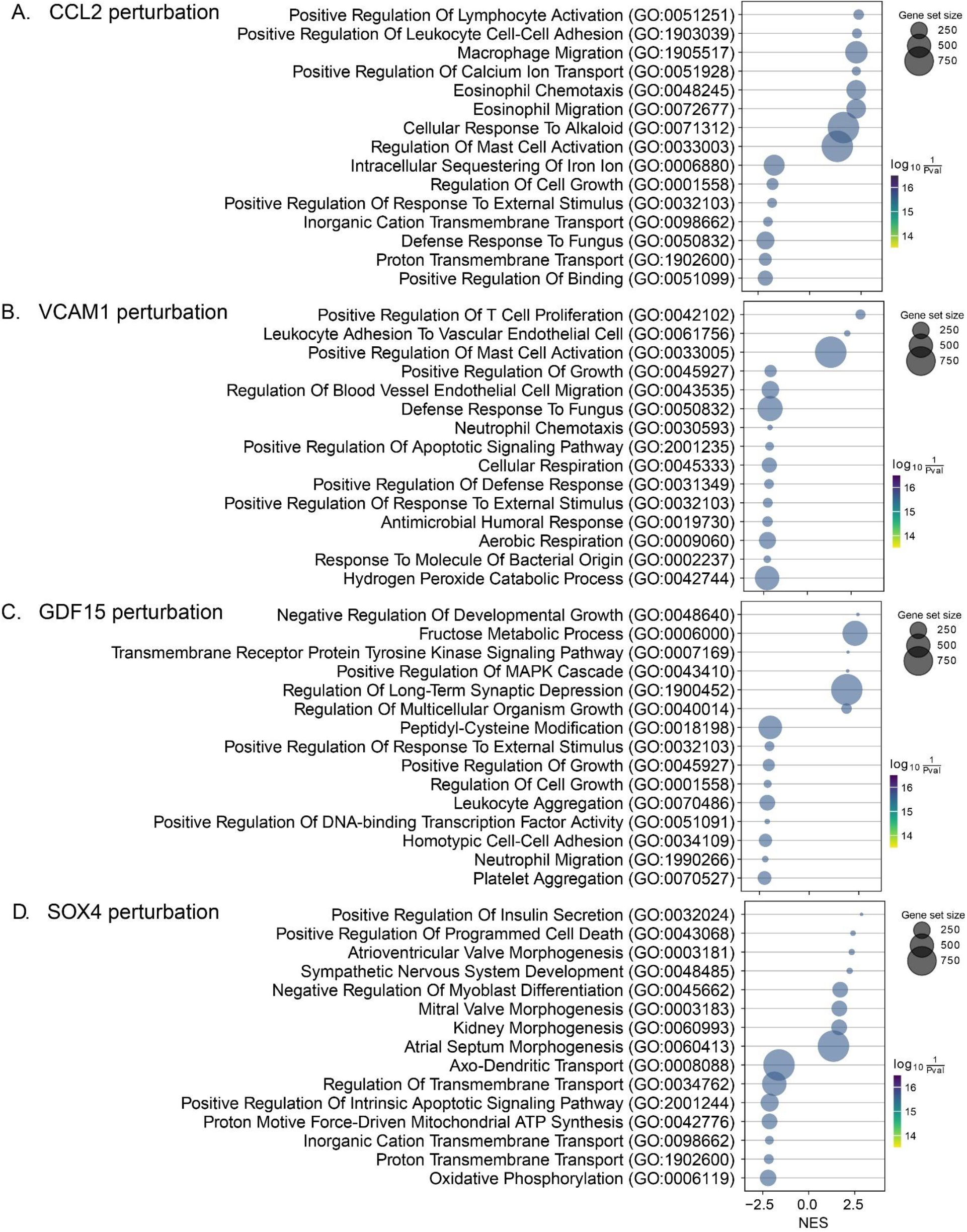
Expression-driven Gene Ontology (GO) enrichment results for four gene perturbations. Panels A-D show the top enriched GO Biological Process terms following perturbation of CCL2 (A), VCAM1 (B), GDF15 (C) and SOX4 (D). Each bubble represents one GO term; the x-axis shows the normalized enrichment score (NES), bubble size corresponds to the gene set size (number of genes in the GO term), and bubble color encodes statistical significance as −log10(p-value) (darker = more significant). Terms are ordered by significance and effect size and only the most enriched / interpretable terms are displayed for clarity. Positive NES values indicate enrichment among up-regulated genes after perturbation, while negative NES values indicate enrichment among down-regulated genes. Enrichment was calculated using gene set enrichment analysis (GSEA) on ranked differential expression results, and GO terms shown are from the Biological Process ontology.

## Discussion

Nephrobase Cell+ represents a new paradigm in kidney genomics by providing a *foundation model* pretrained on an unprecedented scale of kidney data. Its training on ~39.5 million cells and nuclei across four species (human, mouse, rat, pig) and diverse modalities (scRNA-seq, snRNA-seq, spatial transcriptomics, snATAC-seq) equips the model to capture conserved kidney biology while tolerating technical variation. In contrast, conventional tools like Seurat and Scanpy are task-specific frameworks for single assays: they excel at preprocessing, visualization, clustering and simple data integration, but they lack the ability to learn a unified latent representation through generative pretraining. For example, Seurat’s anchoring approach can align datasets across modalities^20^, and Scanpy scales to millions of cells^17^, but neither provides transferable cell or gene embeddings that predict expression patterns or responses to perturbation. Likewise, recent foundation models such as scGPT demonstrate the promise of pretraining on large cell repositories^3^, but by training on multi-organ datasets they may not capture kidney-specific regulatory structure as effectively. Nephrobase Cell+ fills this gap by specializing in kidney tissue: its organ-centric training helps the model learn hierarchical nephron organization, segment-specific pathways, and microenvironmental cues unique to the kidney.

Single-cell “foundation” models like Geneformer and scFoundation have recently been introduced to capture broad transcriptional patterns from large atlas datasets. Geneformer is a transformer encoder pretrained on ~30 million human single-cell transcriptomes^21^. It learns gene-gene relationships in a self-supervised masked-learning framework and, when fine-tuned on limited data, boosts accuracy on diverse network biology tasks (e.g. chromatin state prediction and cell-type classification). Similarly, scFoundation (Hao *et al*., 2024) uses an asymmetric transformer-like masked-autoencoder architecture pretrained on ~50 million human single-cell profiles covering ~20,000 genes^22^. scFoundation achieved state-of-the-art performance on a wide array of single-cell tasks, including imputation of missing gene expression, prediction of drug-response at the tissue and single-cell level, cell type annotation, and perturbation outcome prediction. In contrast, GEARS (Lotfollahi et al., 2023) takes a very different approach: it is not a general “language model” for gene expression, but rather a graph-based perturbation simulator. GEARS embeds each gene and each perturbation as trainable vectors and combines them via a graph neural network based on known gene-gene relationships^23^. In practice GEARS is trained on single-cell perturb-seq screens, learning to predict transcriptional responses to new single-or multi-gene perturbations. Lotfollahi *et al*. ^23^ show that GEARS can forecast combinatorial perturbation outcomes with ~40% higher precision than previous methods and can uncover complex genetic interactions even for genes never jointly perturbed in the data.

By integrating multi-modal and cross-species data, Nephrobase Cell+ addresses key challenges in nephrology. Chronic kidney disease (CKD) is a major global health burden^24^, yet its mechanisms are complex, involving interactions between epithelial, endothelial, immune, and stromal cells. Single-cell and spatial studies have begun to map cell states and microenvironments in kidney disease ^25, 26^, but batch effects and sparse coverage complicate analysis. Nephrobase Cell+’s adversarial and contrastive training components explicitly remove assay- and batch-specific signals, yielding embeddings that are more *assay-invariant*. For instance, when applied to multi-modal data from healthy and diseased kidneys, the model can project spatial transcriptomic profiles (CosMx, Xenium) and dissociated single-cell data into the same latent space, enabling direct comparison of microenvironment composition. Indeed, recent work using integrated scRNA/snRNA and spatial profiles identified four distinct kidney microenvironments (glomerular, immune, tubular, fibrotic) that correlate with disease state^25^; Nephrobase Cell+ can generalize this approach, providing a unified mapping from any input assay to a spatially-informed kidney atlas. Such integration unlocks new insights: for example, Nephrobase Cell+ could reveal how pathogenic fibroblast or immune niches emerge in CKD and identify conserved gene programs driving fibrosis across samples.

Compared to established computational pipelines, Nephrobase Cell+ offers advantages in scalability, accuracy, and generalizability. Its transformer architecture is designed to handle large gene sets and can be scaled up (here to ~1 billion parameters) much as NLP models have been, allowing it to absorb vast training data^3^. The model’s performance benchmarks on held-out kidney data far exceed those of simpler methods: its higher NMI/ARI indicates more accurate cell-type separation than traditional clustering, and its favorable batch-correction metrics (iLISI, kBET) demonstrate robust integration even across species and protocols. Moreover, by learning gene-gene dependencies through attention, Nephrobase Cell+ may implicitly capture regulatory networks that are not easily accessible to tools like Scanpy or Seurat. Importantly, the learned embeddings are general-purpose: once trained, Nephrobase Cell+ can be fine-tuned for downstream tasks (e.g. cell-type annotation, trajectory inference, or simulation of perturbation responses) with minimal additional data. This adaptability goes beyond what static tools provide; for example, scGPT has shown that foundational representations can be transferred to tasks like batch integration and perturbation prediction, and we anticipate Nephrobase Cell+ will similarly accelerate nephrology applications by providing pretrained features tailored to kidney biology.

Nonetheless, there are limitations. Although extensive, the Nephrobase Cell+ training set does not cover every possible kidney condition or assay. Some rare cell types or extreme pathologic states may still be underrepresented, potentially limiting the model’s performance on those instances. The choice of a fixed gene feature space (32,768 orthologous genes) means genes outside this set cannot be directly handled. Like all large models, Nephrobase Cell+ requires substantial computational resources to train, which may restrict re-training or extension by typical labs. In its current form, Nephrobase Cell+ models primarily RNA and ATAC modalities; other data types (e.g. proteomics, metabolomics, imaging-derived phenotypes) and species (non-mammalian models) are not yet integrated. Finally, while foundation models can capture correlations, they do not alone prove causation; experimental validation remains essential for any new hypothesis generated.

Looking forward, Nephrobase Cell+ opens many research directions. Future work should expand the model to include additional species (e.g. non-human primates) and richer modalities (spatial multi-omics, proteogenomic data) as those datasets emerge. Fine-tuning Nephrobase Cell+ on specific CKD subcohorts or organoid models could improve diagnostic classification or drug response prediction in nephrology. The attention weights and latent spaces learned by the model could be mined to discover novel regulatory circuits or to prioritize candidate biomarkers across cell types. Finally, iterative updating of the model with new data - including patient-derived biopsy data and clinical outcomes - could help bridge the gap between molecular signatures and patient prognosis. In summary, Nephrobase Cell+ lays a versatile foundation for kidney research, combining the strengths of massive data integration with the flexibility of deep learning. By overcoming fragmentation and scale challenges in nephrology data, it is poised to drive new insights into CKD mechanisms, kidney cell heterogeneity, and microenvironmental pathobiology that were previously out of reach.

## Method

### Data Acquisition

Our dataset was assembled to create a comprehensive multi-species, multi-modal atlas of kidney biology, totaling approximately 40 million single-cell or single-nucleus profiles. This dataset spans four mammalian species: human (*Homo sapiens*), mouse (*Mus musculus*), rat (*Rattus norvegicus*), and pig (*Sus scrofa*). It encompasses various relevant biological contexts, including adult kidney tissue, fetal kidney development, kidney organoids, and peripheral blood mononuclear cells derived from both healthy donors and individuals diagnosed with Chronic Kidney Disease (CKD). The data integrates extensive publicly available resources with substantial internally generated datasets. Public data were systematically curated from major repositories such as the Gene Expression Omnibus (GEO)^27^, Sequence Read Archive (SRA), Human Cell Atlas (HCA)^28^, the CELLxGENE database^29^, the Kidney Precision Medicine Project (KPMP)^8^, and other relevant consortia outputs, filtering for the target species and biological samples. In addition to public data, we generated substantial multi-modal data in-house to enhance the dataset’s diversity. This includes ~3 million cells profiled using CosMx^25^ Spatial Molecular Imager (NanoString) and ~2 million cells using Xenium^30^ In Situ (10x Genomics), providing high-plex spatial transcriptomic information. Furthermore, we generated ~3 million single-nucleus, single-cell and single-nucleus assay for transposase-accessible chromatin using sequencing (snATAC-seq).

### Gene Orthology Mapping and Feature Space Harmonization

To enable cross-species analysis, gene identifiers from mouse, rat, and pig datasets were mapped to their human orthologs using annotations from Ensembl^31^ release 113. We prioritized high-confidence, one-to-one orthology relationships. Based on this mapping and potentially considering gene variance or representation across datasets, a final unified feature space comprising exactly top 32,768 highly variable genes were selected for model training. This space primarily utilizes human gene symbols corresponding to ortholog groups, allowing the model to leverage conserved biological information while species-specific context was provided through dedicated input embeddings.

### In-house Sample Acquisition

The University of Pennsylvania institutional review board (IRB) approved the collection of human kidney tissue for this study. Left-over kidney samples were irreversibly de-identified, and no personal identifiers were gathered. Therefore, they were exempt from IRB review (category 4). We engaged an external, honest broker responsible for clinical data collection without disclosing personally identifiable information. Participants were not compensated.

### snRNA-seq

Nuclei were isolated using lysis buffer (Tris-HCl, NaCl, MgCl_2_, NP40 10% and RNAse inhibitor (40 U μl^−1^)). In total, 10-30 mg of frozen kidney tissue was minced with a razor blade into 1-2 mm pieces in 1 ml of lysis buffer. The chopped tissue was transferred into a gentleMACS C tube and homogenized in 2 ml of lysis buffer using a gentleMACS homogenizer with programs of Multi_E_01 and Multi_E_02 for 45 s. The homogenized tissue was filtered through a 40 µm strainer (Thermo Fisher Scientific, 08-771-1), and the strainer was washed with 4 ml wash buffer. Nuclei were centrifuged at 500*g* for 5 min at 4 °C. The pellet was resuspended in wash buffer (PBS 1× + BSA 10% (50 mg ml^−1^) + RNAse inhibitor (40 U μl^−1^)) and filtered through a 40 µm Flowmi cell strainer (Sigma-Aldrich, BAH136800040-50EA). Nuclear quality was checked, and nuclei were counted. In accordance with the manufacturer’s instructions, 30,000 cells were loaded into the Chromium Controller (10X Genomics, PN-120223) on a Chromium Next GEM Chip G Single Cell Kit (10X Genomics, PN-1000120) to generate single-cell GEM (10X Genomics, PN-1000121). The Chromium Next GEM Single Cell 3′ GEM Kit v3.1 (10X Genomics, PN-1000121) and Single Index Kit T Set A (10X Genomics, PN-120262) were used in accordance with the manufacturer’s instructions to create the cDNA and library. Libraries were subjected to quality control using the Agilent Bioanalyzer High Sensitivity DNA Kit (Agilent Technologies, 5067-4626). Libraries were sequenced using the NovaSeq 6000 system (Illumina) with 2 × 150 paired-end kits. Demultiplexing was as follows: 28 bp Read1 for cell barcode and UMI, 8 bp I7 index for sample index and 91 bp Read2 for transcript.

### snATAC-seq

The procedure described above for snRNA-seq was used to isolate the nuclei for ATAC-seq. The resuspension was performed in diluted nuclei buffer (10× Genomics). Nuclei quality and concentration were measured in the Countess AutoCounter (Invitrogen, C10227). Diluted nuclei were loaded and incubated in chromium single-cell ATAC Library and Gel Bead Kit’s transposition mix (10X Genomics, PN-1000110). Chromium Chip E (10X Genomics, PN-1000082) in the chromium controller was used to capture the gel beads in the emulsions (GEMs). The Chromium Single Cell ATAC Library & Gel Bead Kit and Chromium i7 Multiplex Kit N Set A (10X Genomics, PN-1000084) were then used to create snATAC libraries in accordance with the manufacturer’s instructions. Library quality was examined using an Agilent Bioanalyzer High Sensitivity DNA Kit. After sequencing on an Illumina Novaseq system using two 50 bp paired-end kits, libraries were demultiplexed as follows: 50 bp Read1 for DNA fragments, 8 bp i7 index for sample index, 16 bp i5 index for cell barcodes and 50 bp Read2 for DNA fragments.

### scRNA-seq

Fresh human kidneys (0.5 g) collected in RPMI media were minced into approximately 2-4 mm cubes using a razor blade. The minced tissue was then transferred to a gentleMACS C tube containing Multi Tissue Dissociation Kit 1 (Miltenyi Biotec, 130-110-201). The tissue was homogenized using the Multi_B program of the gentleMACS dissociator. The tube, containing 100 μl of enzyme D, 50 μl of enzyme R and 12.5 μl of enzyme A in 2.35 ml of RPMI, was incubated for 30 min at 37 °C. Second homogenization was performed using the Multi_B program on the gentleMACS dissociator. The solution was then passed through a 70-μm cell strainer. After centrifugation at 600*g* for 7 min, the cell pellet was incubated with 1 ml of RBC lysis buffer on ice for 3 min. The reaction was stopped by adding 10 ml of PBS. Next, the solution was centrifuged at 500*g* for 5 min. Finally, after removing the supernatant, the pellet was resuspended in PBS. Cell number and viability were analyzed using Countess AutoCounter (Invitrogen, C10227). This method generated a single-cell suspension with greater than 80% viability. Next, 30,000 cells were loaded into the Chromium Controller (10X Genomics, PN-120223) on a Chromium Next GEM Chip G Single-Cell Kit (10X Genomics, PN-1000120) to generate single-cell GEM according to the manufacturer’s protocol (10X Genomics, PN-1000121). The cDNA and library were made using the Chromium Next GEM Single Cell 3′ GEM Kit v3.1 (10X Genomics, PN-1000121) and Single Index Kit T Set A (10X Genomics, PN-120262) according to the manufacturer’s protocol. Quality control for the libraries was performed using the Agilent Bioanalyzer High Sensitivity DNA Kit (Agilent Technologies, 5067-4626). Libraries were sequenced on the NovaSeq 6000 system (Illumina) with 2 × 150 paired-end kits using the following demultiplexing: 28 bp Read1 for cell barcode and unique molecular identifier (UMI), 8 bp I7 index for sample index and 91 bp Read2 for transcript.

### Single Nuclei and Cell RNAseq Data Processing

FASTQ files from each 10X single nuclei/cell run were processed using Cell Ranger v9.0.1 (10X Genomics). Gene expression matrices for each cell were produced using the human genome reference GRCh38 or GRCh37, mouse genome reference GRCm39, rat genome reference mRatBN7.2, Sus scrofa genome reference Sscrofa11.1. Ambient RNA was corrected using CellBender^32^. Initial quality control involved filtering cells with fewer than 200 unique molecular identifiers to remove low-quality cells. To identify and remove outlier cells based on quality control metrics, we employed a median absolute deviation (MAD) approach. Cells were flagged as outliers if their log-transformed total counts, log-transformed number of genes detected, or percentage of reads in the top 20 genes fell outside of a range defined by ±5 MADs from the median for each respective metric. Finally, to remove genes with extremely low expression across the dataset, we filtered out genes that were detected in fewer than one cell. This multi-step filtering process resulted in a refined dataset suitable for downstream analyses.

### Single Nuclei ATACseq Data Processing

Raw FASTQ files were aligned to GRCh38 and quantified via Cell Ranger ATAC (v1.1.0). Low-quality cells were filtered (criteria: peak_region_fragments <3000 & >20000, pct_reads_in_peaks <15, nucleosome_signal >4, TSS.enrichment <2). Filtered cells were merged in Seurat. Dimension reduction involved SVD of the TFIDF matrix and UMAP.

### CosMx Sample preparation and data preprocessing

Tissue sections were cut at 5 µm thickness and prepared according to the manufacturer specifications (NanoString Technologies). We used the human universal cell characterization RNA probes, and 50 additional custom probes for the following genes: ESRRB, SLC12A1, UMOD, CD247, SLC8A1, SNTG1, SLC12A3, TRPM6, ACSL4, SCN2A, SATB2, STOX2, EMCN, MEIS2, SEMA3A, PLVAP, NEGR1, SERPINE1, CSMD1, SLC26A7, SLC22A7, SLC4A9, SLC26A4, CREB5, HAVCR1, REN, AP1S3, LAMA3, NOS1, PAPPA2, SYNPO2, RET, LHX1, SIX2, CITED1, WNT9B, AQP2, SCNN1G, ALDH1A2, CFH, NTRK3, WT1, NPHS2, PTPRQ, CUBN, LRP2, SLC13A3, ACSM2B, SLC4A4, PARD3, XIST, UTY. We used DAPI, CD298/B2M, CK8/18, and PanCK/CD45 for additional staining per the Nanostring protocol. Imaging was performed using configuration A. After imaging was completed, the flowcell was incubated in 100% xylene overnight, the coverslip was removed from the slide with a razor blade, and the slide was then stained with hematoxylin and eosin. The expression matrix and metadata from each CosMx run were exported from the AtoMx platform and converted to a Python object using Squidpy. All samples were merged, preprocessed, and analyzed together using Scanpy. Cells with fewer than 30 counts were filtered out.

### Xenium Sample preparation and data preprocessing

Tissue sections were cut at 5 µm thickness and cut onto a Xenium slide according to the manufacturer specifications (10X Genomics). We used the human Xenium Prime 5K Human Pan Tissue & Pathways Panel with 100 additional custom probes for the following genes: TPM1, ESRRB, COL6A3, AGR2, SLC26A7, ATP1B1, SLC8A1, ATP6AP2, TAGLN, SPP1, SAT1, MYL9, LDB2, DEFB1, COL1A2, ACTA2, ST6GALNAC3, SLC13A3, SLC12A3, SLC12A1, MGP, IGHG1, FN1, C7, ACSM2B, AIF1, APOE, AQP3, AZGP1, C1QA, C1QB, C1QC, CAV1, PPIA, CD74, CHI3L1, COL1A1, COL6A1, CRYAB, CXCL14, ENO1, HLA-DPA1, HLA-DRA, IFI27, IGHA1, IL32, KLF2, LGALS3, LUM, MMP7, PIGR, S100A2, SLC4A4, SLPI, SOD2, SPINK1, SOX4, SPOCK2, TACSTD2, TM4SF1, TPM2, VIM, ZFP36, AQP2, RNASE1, ALDOB, PCGF6, RHOB, CD81, ASS1, MYL6, COX8A, CTSB, GATM, MT1G, TMSB10, COL3A1, MIF, TPT1, COL6A2, BST2, CLU, APOC1, APOD, PHKG2, RGCC, HLA-DQA2, CORO1A, HSPB1, ADIRF, CKB, HLA-DQB1, COX5B, MT1H, RAMP3, TYROBP, LAMTOR5, ITM2B, UBB, CTSD. Additionally, the same sections were stained according to the Xenium Cell Segmentation workflow for automated morphology-based cell segmentation, and subsequently loaded onto the Xenium Analyzer for in situ transcriptomic analysis. Xenium raw output files were processed using the spatialdata framework (v0.0.14) with the spatialdata-io Xenium plugin. Xenium transcript and segmentation data were loaded from the manufacturer’s output directory using the xenium() function, which parses transcript tables, cell segmentation boundaries, and spatial metadata into a structured SpatialData object. The gene expression table was extracted as an AnnData object for downstream single-cell analysis. Cells with fewer than 30 counts were filtered out.

### Nephrobase Cell+

Our model, Nephrobase Cell+, is designed for single-cell gene expression analysis and cell type classification. It employs Transformer-based encoder-decoder architecture with specialized modules for gene and numerical feature embedding, mixture of experts’ layers, and optional adversarial domain/assay adaptation.

### Gene Encoding

We represent each gene as a unique index and employ a trainable embedding layer to map these indices into a continuous vector space. Let *G* be the number of genes, and *d*_*embed*_ be the embedding dimension. The gene embedding layer, *E*_*gene*_, is a matrix of size *G* × *d*_*embed*_. For a gene index input *g* ∈ {0,1, …, *G* − 1}, the gene embedding **e**_*g*_ is obtained by:**e**_*g*_ = *E*_*gene*_[*g*], where *E*_*gene*_[*g*] denotes the *g*-th row of the embedding matrix *E*_*gene*_. The output 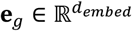 represents the embedded vector for gene *g*.

### Gene Expression Encoding

In addition to gene indices, our model incorporates numerical features derived from gene expression counts to provide richer input representation. To effectively use gene expression counts as numerical features, we first preprocess the raw count data using sum-log normalization to account for variations in sequencing depth and stabilize variance inherent in count data.

#### Sum-Log Normalization of Gene Counts

Prior to being fed into the numerical feature embedding module, raw gene counts undergo sum-log normalization. For each sample *i* and gene *j* in the input count matrix *X* ∈ ℝ^*B*×*G*^, where *B* is the batch size and *G* is the number of genes, we calculate the normalized and transformed count *x*′_*ij*_ using the formula: 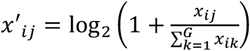. This process yields a matrix *X*′ ∈ ℝ^*B*×*G*^ of sum-log normalized gene expression values.

#### Embedding Normalized Gene Expression

For each gene *j*, we treat its sum-log normalized expression value (which is a single numerical value *x*′_*ij*_ for each sample *i* in a batch) as the numerical feature to be embedded. In a typical scenario where we are processing gene features independently, and assuming we are embedding a single representative numerical value for each gene (or potentially processing each sample’s normalized count for each gene separately and then aggregating - clarification needed on the exact input to the embedding layer in the broader model context if it’s not a single value), we can consider the input to the embedding layer as a numerical feature *x* ∈ ℝ^*size*^, where in the simplest case, *size* = 1, representing a single, normalized gene expression value. The embedding process for this numerical feature *x* then involves a series of linear transformations, a non-linear activation function, and a dropout layer. Let *d*_*embed*_ denote the embedding dimension and let *mlp*_*ratio* control the width of hidden layers within the embedding network. The numerical embedding process can be described as follows:

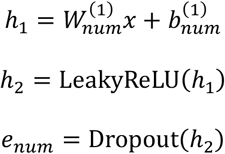

where 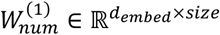 and 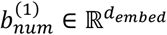 are the weights and bias of the linear layer, respectively. LeakyReLU represents the Leaky ReLU activation function, and Dropout denotes the dropout operation. The output 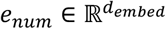 is the resulting embedded vector for the numerical feature *x*, representing the learned embedding of the gene’s expression information.

### Root Mean Square Layer Normalization (RMSNorm)

We use RMSNorm for stabilization. RMSNorm normalizes the input tensor *x* based on its root mean square^33, 34^.

### Multi-Layer Perceptron (MLP)

Non-linear transformations are performed using an MLP layer. Our MLP consists of three linear layers (*w*_1_, *w*_2_, *w*_3_) and a SiLU activation gate^35, 36^. The forward pass is computed as:

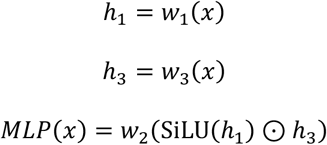

where SiLU is the Sigmoid Linear Unit activation, and ⊙ denotes element-wise multiplication.

### Mixture of Experts (MoE)

Our MoE layer follows the sparse Top-k routing paradigm^37^, where experts are dynamically selected based on SoftMax probabilities and combined via weighted summation^38^. This design aligns with scalable MoE architectures validated in large language models^39^. For an input feature vector 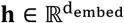, the routing process is as follows: a) Router Logits: A linear layer, 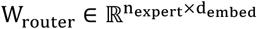 and 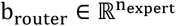, calculates logits for each expert: l = W_router_h + b_router_ where 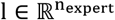, and n_expert_ is the number of experts. b) Routing Probabilities: The logits are converted into probabilities using a SoftMax function: p = softmax(l), where 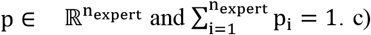 Expert Selection: The top-k experts with the highest probabilities are selected. Let I_topk_ be the indices of the top-k experts. d) Expert Weights: The probabilities of the selected experts are normalized to sum to 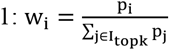 for i ∈ I_topk_. e) Expert Computation and Combination: Each selected expert, E_i_ (implemented as a basic MLP module), processes the input h. The final output o is a weighted sum of the outputs from the selected experts: 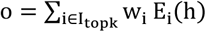.

#### Shared MoE

The Shared MoE module extends the MoE by adding a set of shared experts^40^. The final output is the sum of the outputs from the MoE and the shared experts. If *S*_*j*_ represents the *j*-th shared expert, and *n*_*shared*_ is the number of shared experts, the output o_*shared*_*MOE*_ of the Shared MoE for input h is: 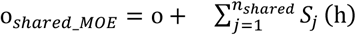, where o is the output from the MoE component.

#### Load Balancing Loss for MoE

To encourage balanced expert utilization in MoE, we incorporate a load balancing loss, *L*_*load*_*balance*_.^38^ This loss aims to ensure that experts are used more uniformly during training. Let 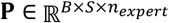 be the router probabilities for a batch of *B* sequences of length *S*. The load balancing loss is calculated as: *L*_*load*_*balance*_ = aux_loss + z_loss × z_loss_weight, where:

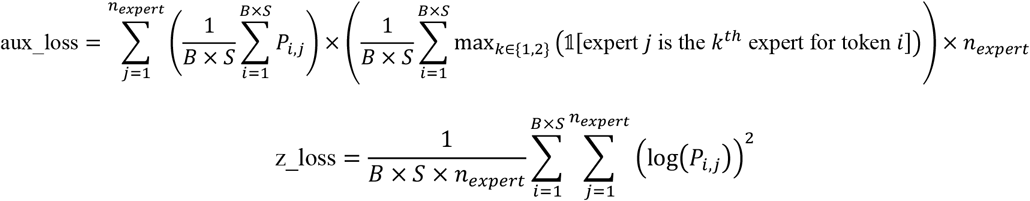

and z_loss_weight is a small weight (e.g., 0.001).

### Elastic cell similarity (ECS)

ECS loss serves as a regularization term that encourages cell embeddings^3^ to be dissimilar from each other, up to a certain threshold. This promotes diversity in the embedding space and can prevent collapse, where all cells are mapped to similar representations. The ECS loss is calculated as follows:

Given a tensor of cell embeddings, *E* = [*e*_1_, *e*_2_, …, *e*_*n*_], where *e*_*i*_ is the embedding for the *i*-th cell and *n* is the number of cells in the batch, we first normalize each embedding vector to unit length: 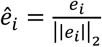 where ||*e*_*i*_ ||_2_ is the L2 norm of *e*_*i*_. Let Ê = [Ê_1_, Ê_2_, …, Ê_*n*_] be the matrix of normalized embeddings. We compute the cosine similarity matrix *C* between all pairs of normalized embeddings. The element *C*_*ij*_ of this matrix represents the cosine similarity between the *i*-th and *j*-th embedding: 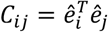. This can be efficiently computed using matrix multiplication: *C* = ÊÊ^*T*^. To avoid comparing an embedding with itself, we mask the diagonal elements of the cosine similarity matrix. Then, we calculate the ECS loss, *L*_*ECS*_, as the mean squared error between the off-diagonal elements of the cosine similarity matrix and a predefined threshold τ_*ecs*_:

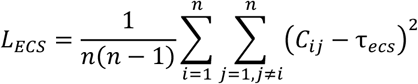

This can be implemented by first setting the diagonal of *C* to zero, and then calculating the mean of the squared differences: *L*_*ECS*_ = Mean((*C* − τ_*ecs*_1)^2^ ⊙ (1 − *I*)), Where 1 is a matrix of ones with the same dimensions as *C, I* is the identity matrix, and ⊙ denotes element-wise multiplication. The threshold τ_*ecs*_ is a hyperparameter, typically set to a value like 0.5, controlling the desired level of dissimilarity.

### Supervised Contrastive Loss

The Supervised Contrastive Loss is employed when label information is available^41-43^. It aims to pull embeddings of samples with the same label closer together while pushing embeddings of samples with different labels further apart. Similar to ECS, the input embeddings are first normalized: 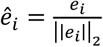. A similarity matrix *S* is computed using the normalized embeddings and a temperature parameter 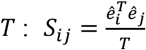 where *T* is a temperature scaling factor, typically a small positive value (e.g., 0.07). For each sample *i*, we identify samples that have the same label. A binary mask matrix *M* is created where *M*_*ij*_ = 1 if sample *i* and sample *j* have the same label (and *i* ≠ *j*), and *M*_*ij*_ = 0 otherwise. Formally, if *l*_*i*_ is the label of sample *i*, then:

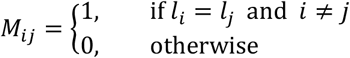

For each sample *i*, we want to maximize the similarity with positive samples (samples with the same label) and minimize the similarity with negative samples (samples with different labels). The loss for each sample *i* is defined based on the log-softmax of the similarities, focusing on positive pairs:

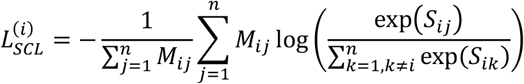

This formula calculates the negative log-likelihood of correctly classifying the positive samples among all other samples. The term 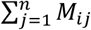 is the count of positive samples for sample *i*. The overall Supervised Contrastive Loss is the average over all samples: 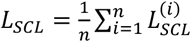.

### Loss Function for Zero-Inflated Negative Binomial (ZINB) Regression

To model count data exhibiting overdispersion and zero-inflation, we employed a ZINB regression loss function^3, 22, 44, 45^. This loss function is particularly suited for scenarios where observed counts are derived from a mixture of two processes: one generating counts from a Negative Binomial (NB) distribution and another process generating excess zeros. The ZINB distribution is parameterized by a mean parameter (*μ*), a dispersion parameter (*θ*), and a zero-inflation probability (*π*). The ZINB probability mass function for a count *y* is defined as:

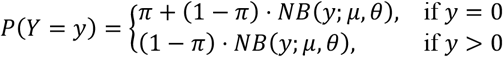

where *NB*(*y*; *μ, θ*) represents the probability mass function of the Negative Binomial distribution, parameterized by mean *μ* and dispersion *θ*. Specifically, we parameterize the Negative Binomial distribution in terms of mean and dispersion, where the variance is given by 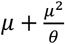.

The negative log-likelihood (NLL) loss for the ZINB model, which we aim to minimize, is derived from this probability mass function. For a given observation *y*_*i*_, predicted mean *μ*_*i*_, predicted dispersion *θ*_*i*_, and predicted zero-inflation probability *π*_*i*_, the ZINB loss (ℒ_*ZINB*_) is formulated as:

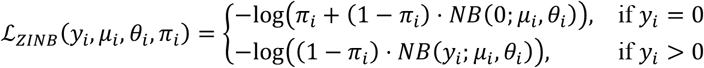

In practice, to ensure numerical stability and differentiability, we implemented the loss using softplus and log-gamma functions. The Negative Binomial log-likelihood component, *NB*(*y*; *μ, θ*), was calculated as:

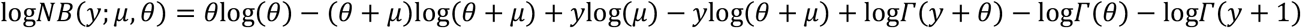

where *Γ*(⋅) is the gamma function. To further enhance numerical stability and handle the zero-inflation probability *π*, we utilized the softplus function, *softplus*(*x*) = log(1 + *e*^*x*^), and parameterized the zero-inflation component using logits (*ρ*) such that *π* = sigmoid 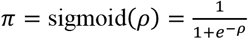. In our implementation, we directly predicted the zero-inflation logits (*ρ*), denoted as zero_logits in our model outputs. The total loss for a batch of observations was computed as the means of the individual ZINB losses across all data points in the batch.

Prior to applying the ZINB loss, we performed total count normalization on the input count data. For each sample, we calculated the sum of all counts and scaled each count such that the total sum for each sample was normalized to a target value of 10^4^. This normalization step, implemented as: 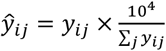, where *y*_ij_ is the original count for feature *j* in sample *i*, and ŷ_*ij*_ is the normalized count. This step mitigates the effect of varying sequencing depths across samples, ensuring fair comparison and model training.

### Adversarial Network

To eliminate assay-specific and batch-specific biases in feature representations, we integrate an adversarial training framework. This framework employs a MLP discriminator and a gradient reversal layer (GRL) ^46^ to adversarially optimize the feature generator. The discriminator is trained to classify assay/batch labels from the input features, while the generator learns to confound these predictions via GRL-based gradient inversion. A dynamic loss scaling strategy further refines the adversarial objective, prioritizing bias removal as training progresses. This dual adversarial mechanism ensures robust, assay/batch-invariant representations for downstream tasks^47^.

#### Adversarial Discriminator Architecture

We employed a MLP as the adversarial discriminator, denoted as *D*. This discriminator network is designed to classify the domain or assay of the input feature representations. The discriminator *D* consists of *n*_*layers*_ layers. The discriminator *D*(*h*) is computed through a series of transformations. Let *h* be the input feature representation, *h*_1_ = *W*_1_*h* + *b*_1_.

For *i* = 2,3, …, *n*_*layers*_:

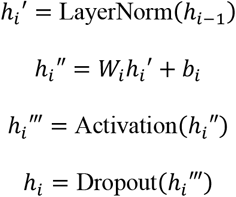

Finally, the output layer is:

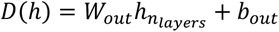

Where *W*_*i*_ and *b*_*i*_ are the weights and biases of the *i*-th linear layer, respectively. LayerNorm represents Layer Normalization, Activation is a non-linear activation function (LeakyReLU), and Dropout is applied with a probability of 0.3. The dimensions of the weight matrices are configured to achieve a hidden dimension of *d*_*model*_ × *mlp*_*ratio*. The final linear layer projects to *n*_*cls*_ output classes, where *n*_*cls*_ represents the number of domains or assays to be discriminated against.

#### Gradient Reversal Layer

To facilitate adversarial training, a GRL was inserted before the input to the discriminator. The GRL acts as an identity function during the forward pass but reverses the gradient by multiplying it by −*λ* during backpropagation. Formally, for an input *x*, the GRL operation *GRL*(*x*) and its gradient behavior are defined as:

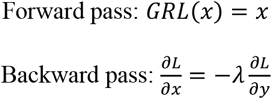

where *y* = *GRL*(*x*) and 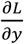 is the gradient from subsequent layers. *λ* is a hyperparameter controlling the strength of gradient reversal.

#### Adversarial Loss Functions

Cross-entropy loss is used as the objective function for both domain and assay adversarial tasks. For domain adversarial training, the objective is to minimize the discriminator’s ability to correctly identify the domain, thus encouraging domain-invariant feature learning in the main network. The adversarial domain loss *L*_*adv*_*domain*_ is defined as: *L*_*adv*_*domain*_ = *L*_*CE*_(*D*_*domain*_(*GRL*(*h*)), *y*_*domain*_), where *L*_*CE*_ is the cross-entropy loss function, *D*_*domain*_ is the domain discriminator, *h* is the feature representation, and *y*_*domain*_ represents the domain labels. This loss is scaled by a factor *α*_*adv*_*domain*_ to adjust its contribution to the total loss.

For assay adversarial training, the goal is to remove assay-specific biases from the feature representations. The adversarial assay loss *L*_*adv*_*assay*_ is defined similarly: *L*_*adv*_*assay*_ = *L*_*CE*_ (*D*_*assay*_(*GRL*(*h*)), *y*_*assay*_), where *D*_*assay*_ is the assay discriminator and *y*_*assay*_ represents the assay labels. This loss is scaled by *α*_*adv*_*assay*_ and a dynamic scaling factor *s*_*epoch*_ that varies with the training epoch. The dynamic scaling factor *s*_*epoch*_ is defined as:

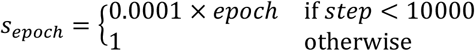

This epoch-dependent scaling progressively increases the influence of the assay adversarial loss during training. The total loss function *L*_*total*_ is a weighted sum of the primary task loss *L*_*main*_, the adversarial domain loss, and the adversarial assay loss:

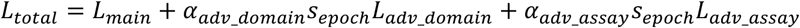

By minimizing *L*_*total*_, the model is trained to learn feature representations that are effective for the primary task while simultaneously being invariant to domain and assay variations, enhancing the model’s generalization capability and robustness.

### Class Imbalance Adjustment

To counteract potential bias arising from class imbalance in the training data, we implemented a class-balanced weighting scheme based on the effective number of samples^48^. Let *n*_*c*_ denote the number of training samples for class *c*. The weight *w*_*c*_ assigned to each class was calculated as: 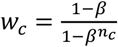, where the hyperparameter *β* was set to 0.9, following ref. 1. This approach assigns higher weights to classes with fewer samples.

The computed weights were subsequently normalized to ensure their mean is unity: 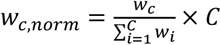, where *C* is the total number of classes. These normalized weights *w*_*c,norm*_ were then used to scale the contribution of each class to the loss function during model training.

### Classification Loss

To address class imbalance and prioritize learning from challenging examples, we utilized the Focal Loss function^1^ as the training objective. Focal Loss^49^ adapts the standard cross-entropy loss by incorporating a modulating factor based on the predicted probability of the true class.

Given the raw output logits **z** from the model for a sample, we first compute the vector of probabilities *p* = softmax(**z**). Let *p*_*t*_ be the predicted probability for the ground-truth class *t*. The Focal Loss (FL) is defined as:

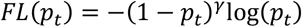

where *γ* is the focusing parameter, set to *γ* = 2.0 in our study. This formulation down-weights the loss contribution from easily classified samples (where *p*_*t*_ is high), thereby increasing the relative importance of misclassified or low-confidence samples.

Computationally, we applied the log-softmax function to the input logits to obtain log-probabilities. The log-probability corresponding to the target class, log(*p*_*t*_), was then selected based on the target class index. The probability *p*_*t*_ was recovered via exponentiation.

To further account for class frequencies, we incorporated an alpha-weighting factor, *α*_*t*_ : *FL*(*p*_*t*_) = −*α*_*t*_(1 − *p*_*t*_)^*γ*^log(*p*_*t*_). The *α*_*t*_ values used were the normalized class weights derived from the effective number of samples strategy (detailed previously). For each sample in a batch, the appropriate *α*_*t*_ weight corresponding to its ground-truth class was applied. The final loss value for a training batch was computed as the arithmetic mean of the individual focal loss values across all samples within that batch.

### Gene Expression Loss Function

The core of the reconstruction loss is the GX_loss function, denoted as *L*_*GX*_. This function, quantifies the difference between the predicted gene expression distribution and the target gene expression. Let 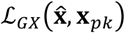 represent the gene expression loss between the model’s output distribution parameters, summarized as 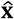, and the target gene expression profile **x**_*pk*_. The specific form of ℒ_*GX*_ is determined by the configuration and may represent various statistical distances or likelihoods depending on the chosen gene expression model (e.g., Zero-Inflated Negative Binomial).

### Loss Calculation

The loss is computed by differentiating between masked and unmasked genes based on a mask *M*_*all*_*flat*_. Let *M*_*all*_*flat*_ be a binary mask indicating which genes are masked. The gene expression loss *L*_*exp*_ is then calculated as a weighted sum of the loss for masked genes and unmasked genes:

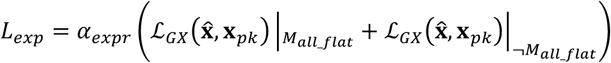

where *α*_*expr*_ is a scaling factor controlling the contribution of the expression reconstruction loss to the total loss. 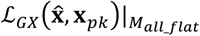 denotes the mean of the gene expression loss evaluated only over the masked genes (where *M*_*all*_*flat*_ is true), and 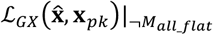 is the mean loss over the unmasked genes (where *M*_*all*_*flat*_ is false). In addition to the gene expression loss *L*_*GX*_, we also monitored the Mean Squared Error (MSE) between the predicted mean expression and the target expression for both masked and unmasked genes as diagnostic metrics, although MSE itself is not directly used as the optimization objective: 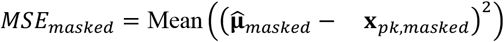 and 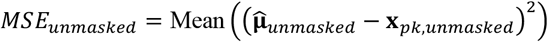, where 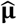 represents the predicted mean expression from the model output, and subscripts *masked* and *unmasked* indicate the regions defined by *M*_*all*_*flat*_.

### Accuracy Metric

To evaluate the performance of the classification task, we computed the classification accuracy. Accuracy is defined as the proportion of correctly classified samples out of the total number of samples.

### Handling of Missing Labels

During training, some samples may have missing or invalid cell type labels, indicated by a label value of −1 in our implementation. To ensure that these samples do not contribute to the classification loss, we filtered out samples with labels equal to −1 before computing the cross-entropy loss and accuracy. Specifically, we only considered samples where *cell_labels* ≠ −1 for loss calculation and accuracy evaluation.

### Loss Scaling

The classification loss was scaled by a factor *α*_*cls*_ to adjust its contribution to the total loss, allowing for fine-tuning the balance between different loss terms if combined with other objectives (e.g., adversarial losses). In our experiments, the classification loss scale was set to 1.0 by default unless otherwise specified.

The minimization of *L*_*cls*_ drives the model to learn feature representations that are discriminative for different cell types, enabling accurate classification of cells based on their learned representations.

### Training Procedure

We employed a Fully Sharded Data Parallel training strategy across 4 H100 GPUs to accelerate the training process. The model was trained end-to-end, minimizing a combined loss function that incorporates both gene expression reconstruction and cell type classification objectives, and optionally adversarial domain and assay adaptation losses.

#### Optimization Algorithm

We used the Adam or AdamW optimizer to update the model parameters. The optimizer was configured with an initial learning rate (*η*), and optionally a weight decay (*λ*) for regularization.

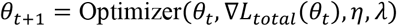

where *θ*_*t*_ represents the model parameters at training step *t*, and *∇L*_*total*_(*θ*_*t*_) is the gradient of the total loss with respect to the parameters.

#### Learning Rate Scheduling

A learning rate scheduler was employed to adjust the learning rate during training. We utilized either a ReduceLROnPlateau scheduler, which reduces the learning rate when validation loss plateaus, or a CosineAnnealingLR scheduler, which follows a cosine annealing schedule. For ReduceLROnPlateau, the learning rate is updated based on validation loss *L*_*val*_:

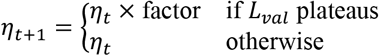

For CosineAnnealingLR, the learning rate follows a cosine function over training steps.

#### Learning Rate Warmup

To stabilize initial training, a linear learning rate warmup strategy was implemented for the first *N*_*warmup*_ steps. During warmup, the learning rate *η*_*t*_ at step *t* is:

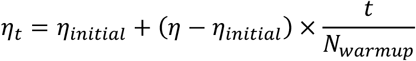

where *η*_*initial*_ is a small initial learning rate (effectively 0 in our setup, starting from a very small value) and *η* is the target learning rate.

#### Gradient Clipping

To prevent exploding gradients, we applied gradient clipping by norm. The gradients were clipped such that their L2 norm does not exceed a predefined threshold (e.g., 1.0).

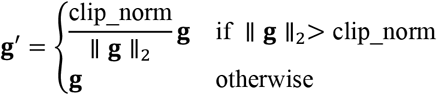

where **g** is the gradient vector, **g**′ is the clipped gradient vector, and clip_norm is the clipping threshold.

#### Mixed Precision Training

To accelerate training and reduce memory consumption, we used Automatic Mixed Precision (AMP) training via torch.cuda.amp.GradScaler and torch.cuda.amp.autocast. This technique performs computations in half-precision (float16) where possible, while maintaining gradients and parameter updates in full precision (float32) for stability.

#### Model Initialization

Model parameters were initialized using Xavier uniform initialization for linear layers, and biases were initialized to zero.

## Data Availability

The previously published data generated for this study are available in GSE107585, GSE182256, GSE183842, GSE173343, GSE211785, GSE209821, GSE183839, and GSE291551. Raw data, processed data, and metadata from the scRNA-seq and CosMx spatial transcriptomics experiments have been deposited in the Gene Expression Omnibus (GEO) under accession code ***, with reviewer token ***.

## Acknowledgments

The authors would like to acknowledge Dylan Jay, David Render, Sean LePeruta, Jovana Lekic, Pio Passariello, and Hannah Hollosi for their valuable contributions and support to this study.

## Authors’ Contribution

Concept and design: CL, MS, NZ and KS; Manuscript drafting: CL, KS; Statistical analysis: CL and EZ; Data collection and interpretation: CL, EZ, BD, JL, EH and SP; Validation: EZ, YS, VR and MS; Manuscript revision: CL, NZ and KS. All authors critically reviewed and agreed to the submission of the final manuscript.

## Funding

This work was supported by the National Institutes of Health grant National Institute of Diabetes and Digestive and Kidney Diseases (NIDDK) R01 DK076077, R01 DK087635, and R01 DK105821 (to KS).

## Notes

### Competing Interest Statement

The authors have declared no competing interest.

